# Nitric oxide synthases from photosynthetic organisms improve growth and confer nitrosative stress tolerance in *E. coli*. Insights on the pterin cofactor

**DOI:** 10.1101/2021.03.07.434255

**Authors:** Natalia Correa-Aragunde, Andrés Nejamkin, Fiorella Del Castello, Noelia Foresi, Lorenzo Lamattina

**Author notes:** Corresponding autor: Lorenzo Lamattina,; Natalia Correa-Aragunde.

## Abstract

Nitric oxide synthase (NOS) synthesizes NO from the substrate L-arginine (Arg). NOS with distinct biochemical properties were characterized from two photosynthetic microorganisms, the unicellular algae *Ostreococcus tauri* (OtNOS) and the cyanobacteria *Synechococcus* PCC 7335 (SyNOS). In this work we studied OtNOS and SyNOS recombinantly expressed in *E. coli* and analyzed bacterial growth and tolerance to nitrosative stress. Results show that the expression of OtNOS and SyNOS promotes bacterial growth and allows metabolizing Arg as N source. In accordance to a high NO producing activity, OtNOS expression induces the *hmp* flavohemoglobin in *E. coli*, suggesting that this strain is sensing nitrosative stress. The addition of 1 mM of the NO donor sodium nitroprusside (SNP) is toxic and generates a strong nitrosative stress. The expression of OtNOS or SyNOS reduced SNP toxicity restoring bacterial growth. Finally, using bioinformatic tools and ligand docking analyses, we propose tetrahydromonapterin (MH4), an endogenous pterin found in *E. coli*, as potential cofactor required for NOS catalytic activity. Our findings could be useful for the development of biotechnological applications using NOS expression to improve growth in bacteria.

**Key points:**

- The NO synthase (NOS) from photosynthetic microorganisms were expressed in *E. coli*
- Expression of NOS increases bacterial growth and tolerance to nitrosative stress.
- Ligand docking analyses indicate tetrahydromonapterin (MH4) as potential NOS cofactor in *E. coli*.

## Introduction

Nitric oxide synthase (NOS) enzymes catalyze the oxidation of L-arginine (Arg) to citrulline and nitric oxide (NO). Animal NOSs are homodimeric enzymes in which each monomer comprises an oxygenase domain and a reductase domain joined by a calmodulin binding motif. The oxygenase domain binds iron protoporphyrin IX (haem), the substrate Arg and the cofactor tetrahydrobiopterin (BH4) at the active site, while reductase domain contains the binding sites of the cofactors NADPH, FMN and FAD (Alderton et al. 2001). In mammals three isoforms were identified, the constitutive endothelial (eNOS) and neuronal NOS (nNOS) and a third inducible NOS (iNOS) found in macrophages (Alderton et al. 2001). The three of them present distinct biochemical properties (Stuehr et al. 2004).

Bacterial NOS, mostly found in Gram positive bacteria, have a single oxygenase domain receiving electrons from promiscuous cellular reductases (Gusarov et al. 2008). NOSs characterized from *Bacillus subtilis, Deinococcus radiodurans* and *Sorangium cellulosum* use tetrahydrofolate (THF) as pterin cofactor since they lack the BH4 synthetic pathway (Pant et al. 2002; Adak et al. 2002b; Reece et al. 2009; Agapie et al. 2009). The biological functions of NOS in bacteria are diverse including tolerance to oxidative/nitrosative stress, secondary metabolite production, and cell growth recovery after oxidative stress (Kers et al. 2004; Gusarov and Nudler 2005; Patel et al. 2009; Zhao et al. 2020).

In the last decade, NOS from photosynthetic organisms were characterized. The NOS from the green alga *Ostreococcus tauri* (OtNOS) is structurally similar to the NOS described in animals (Foresi et al. 2010). Even though, some differences were found in OtNOS like the calmodulin binding domain and the lack of the autoinhibitory loop as occurs in iNOS. Biochemical *in vitro* studies showed that OtNOS oxygenase domain presents an ultrafast NO producing activity. Oxygen activation rate in OtNOS is greater, and NO dissociation was reported 10-80 times faster than in animal NOS (Weisslocker-Schaetzel et al. 2017). OtNOS seems to be constitutively expressed in the alga, although higher expression was observed when the alga was exposed to high light irradiances, suggesting a role in photooxidative stress conditions (Foresi et al. 2010). Lately, the NOS from the cyanobacteria *Synechococcus* PCC 7335 (SyNOS) was identified (Correa-Aragunde et al., 2018). SyNOS architecture has several structural and biochemical features that make it rather different from other NOS described so far, and it challenges the current knowledge about NOSs physiological functions. SyNOS contains an additional globin domain in its N-terminus, followed by the oxygenase and reductase domains (Correa-Aragunde et al. 2018; Picciano and Crane 2019). *In vitro* biochemical analyses indicate that SyNOS globin domain has a NO dioxygenase (NOD) activity, catalyzing the reduction of NO to nitrate (NO_3_^-^) in the presence of O_2_ (Picciano and Crane 2019). SyNOS activity depends on extracellular Ca_2_^+^ (Picciano and Crane 2019), although neither calmodulin nor autoinhibitory loop was found in its structure (Correa-Aragunde et al. 2018). In *in vitro* experiments, it was also reported that SyNOS cannot use THF as cofactor and uses BH4.

Our previous results indicate that the expression of recombinant SyNOS confers *E. coli* the ability to grow at higher OD respect to bacteria expressing the empty vector (EV) and that SyNOS is able to metabolize arginine as N source (Correa-Aragunde et al. 2018). These results suggest that SyNOS may allow bacteria a gain of function to utilize N. However, it remains to confirm which pterin cofactor may SyNOS be using in *E. coli*. Here, we analyze and compare the effect of the expression of OtNOS and SyNOS in *E. coli*. We have studied the growth in different nutrient media, the expression of *hmp* gene coding for the NO dioxygenase in *E. coli* and the tolerance to nitrosative stress. Results indicate that there are some differences between OtNOS and SyNOS regarding *E. coli* growth and nitrosative stress responses. In addition, we have performed *in silico* studies through structural modelling and ligand docking, for contributing to reveal novel insights about the putative pterin factors used by NOS in *E. coli*.

## Materials and Methods

### Bacterial strains and culture conditions

The open reading frames of OtNOS and SyNOS were cloned in the bacterial expression vector pET24a with BamHI-XhOI and EcoRI-XhOI restriction sites respectively. A c-myc tag was added in the C-terminal of SyNOS. The clones were confirmed by DNA plasmid sequencing. Plasmids pET24 (empty vector, EV), pET24a-OtNOS (OtNOS) and pET24a-SyNOS (SyNOS) were used to transform *Escherichia coli* BL21 D3. The expression of recombinant NOS in *E. coli* cultures was induced by addition of 0.1 mM of β-D-1-thiogalactopyranoside (IPTG), 200 μM of aminolevulinic acid (ALA, Sigma) and 1 mM Arginine hydrochloride (Arg, Sigma) in Luria Broth (LB) medium containing 50 μg/ml of kanamycin (Kan, Sigma) and grown at 30 ºC in continuous agitation (200 rpm) to allow oxygenation of the culture. The growth of cultures was followed by OD 600 nm. The detection of recombinant proteins was analyzed by immunoblot using an anti-myc antibody (Sigma) for SyNOS and a specific OtNOS antibody generated by GenScript Company.

### Quantification on nitrate, nitrite and protein content

Bacterial cultures containing Kan 50 μg/ml (OD aprox 0.25) were induced with 0.1 mM IPTG, 200 μM ALA and 1 mM Arg. After 6 h induction, 3 ml of cultures were collected by centrifugation at 5,000 rpm for 5 min and resuspended in 200 μl of 1 X phosphate saline buffer (PBS). Nitrite content was measured by the Griess reagent assay. Fifty μl of samples were incubated for 10 min with 50 μl of 1 % (w/v) sulfanilamide and then 50 μl of 0.1 % (w/v) NED was added. Nitrite content was determined by Abs 540 nm using a solution of sodium nitrite standard curve. Nitrate content was measured by the method described by Cataldo (1975). Briefly, 10 μl of solution was incubated with 5% (w/v) salicylic acid in concentrated H_2_S0_4_ for 20 min. Then the solution was neutralized with 950 μl of 2N NaOH and abs was measured at 401 nm. Protein content in samples was measured by the Bradford method (Bradford 1976).

### RT-PCR analysis

For RT-PCR analysis, *E. coli* cultures were used after 2, 4 and 6 h-induction with 0.1 mM IPTG, 200 μM ALA and 1 mM Arg. Samples were collected by centrifugation at 5,000 rpm for 10 min and RNA was extracted with 1 ml Trizol (Invitrogen) with glass beads. Then, 200 ul of chloroform was added and centrifuged at 10,000 rpm for 15 min. The aqueous solution was isolated, and the RNA precipitated by isopropanol addition and washed by ethanol 70 % (v/v).

The RNA was used for cDNA synthesis in a reaction containing 0.3 μl of random primer, 1 μl of 10 mM dNTP, 2 μl of 0.1 M DTT and 200 U of M-MLV reverse transcriptase (Invitrogen). The *hcaT* gene coding for the MSC transporter (accession number QJZ1029) was used as a reference gene since it is highly invariant in *E. coli* during recombinant protein production in different growth temperatures and induction conditions (Zhou et al. 2011). Primers used for *hmp* were: forward 5’-TCCCTTTACTGGTGGAAACG-3’ and reverse 5’-ACTGTTCCGGTTTGATCTGG-3’, and for the *hcaT* gene were: forward 5’-ATTGCTGCTCGCCAGGGTAG-3’ and reverse 5’-CCAGCGCCATTACCCAGAAC-3’. Real time PCR was performed in StepOne equipment. LinRegPCR program was used for the analysis of qPCR data (Ruijter et al. 2009).

### Growth assay in minimal medium

Bacterial cultures containing 50 μg/ml Kan (OD 600 approximately 0.25-0.3) were induced with 0.1 mM IPTG, 200 μM ALA and 1 mM of Arg for 1.5 h. Aliquots of cultures were diluted (1/100) to 10 ml minimal media containing 5.44 g KH_2_PO_4_ and 6 ml salt solution (10 g MgSO_4_.7H_2_O, 1.0 g MnCl_2_.4H_2_O, 0.4 g FeSO_4_.7H_2_O and 0.1 g CaCl_2_.2H_2_O per l) in 1 l of distilled water. In these growth conditions, the induction of recombinant proteins was maintained by the addition of 0.1 mM IPTG, Kan 50 μg/ml and 200 μM ALA. Filter-sterilized glucose (0.2%, w/v) served as the carbon source with or without filtered-sterilized Arg (0.2%, w/v). The growth was followed by measurement of OD 600 over 30 h at 30 ºC in continuous agitation (200 rpm).

### Evaluation of resistance to nitrosative stress

The evaluation of resistance to nitrosative stress was performed as described by Frey et al. (2002) with minor modifications. *E. coli* Cultures were induced with 0.1 mM IPTG, 200 μM ALA and 1 mM of Arg for 1.5 h in LB medium at 30 ºC and then was used to inoculate 3 ml of fresh LB medium containing either 1 mM nitroprusside sodium dihydrate (SNP; Fluka) to an initial OD 600 of approximately 0.2. Cultures were further grown under identical conditions, and the OD 600 was measured at 30 ºC for 3 h in continuous agitation (200 rpm). Growth rates (μ) were calculated by applying linear regression analysis on the natural logarithm (ln) of Abs 600 values versus time of culture as described by Frey et al. (2002).

### Homology modelling and molecular docking

NOS protein sequences and oxygenase domains were obtained from NCBI database. For tridimensional models building, @TOME v3 Platform (Pons & Labesse - Nucleic Acids Research, Web Server Issue-2009) was used to select the best crystalized template for each NOS of our study. Modeller V9.25 was used for the building of SyNOS model using *Mus musculus* iNOS (PDB code 1M8D) and OtNOS model using *Bos Taurus* eNOS (PDB code 1NSE) as template. Ten models were generated with a DOPE-based loop modeling protocol and the model with the lower DOPE was chosen for docking simulations (Šali and Blundell 1993).

Molecular docking was performed using AutoDock Vina (Trott, O. & Olson. 2010) in Chimera software (Pettersen et al. 2004). First, docking assay was performed for the heam group since it is involved in pterin binding. The binding pocket of the heam group is highly conserved among NOSs and a similar haem bind position was found for OtNOS and SyNOS. Ligands, (6R)-2-amino-5,6,7,8-tetrahydro-6-[(1S,2S)-1,2,3-trihydroxypropyl]pteridin4(3H)-one (MH4) and (6R)-2-amino-6-[(1R,2S)-1,2-dihydroxypropyl]-5,6,7,8-tetrahydro-3H-pteridin-4-one (BH4) were built using Avogadro software (Hanwell et al. 2012) and the energy optimization was determined with PM6 method. Search space for pterins binding poses was set in a grid box of 12×12×12 Å enclosing the arginine and tryptophan necessary for binding. For haem group the search space was set in a grid box of 14×14×14 Å and the position was established taking as reference iNOS superposition. Exhaustiveness was set in 20. In all cases, poses that were not able to bind with the key residues reported were discarded. Visualizations and distance measures were done using Chimera tools.

### Statistical analysis

Results are expressed as means ± standard error. Data were analyzed by one-way analysis of variance (ANOVA) with post hoc Dunnett or Tukey method comparisons. We have developed linear mixed models with normal error structure using the lme function from the nlme library in R software (version 3.1; R Foundation for Statistical Computing). For response variables that do not show a normal distribution, logarithmic transformations were applied, or a generalized linear model has been developed with Gamma error structure in R software. In all cases, transgene expression was set as a fixed effect and experiments were treated as a random effect.

## Results

### Growth of *E. coli* expressing recombinant OtNOS or SyNOS. Analysis of *hmp* expression

We analyzed the effect of OtNOS and SyNOS expression in *E. coli* growth on LB medium. OtNOS and SyNOS expression in *E. coli* was induced by the addition of 0.1 mM IPTG plus 200 uM of the haem precursor aminolevulinic acid (ALA) to a growing culture (OD 0.25) in LB medium containing kanamycin. Bacteria growth was followed by measuring OD 600. Figure 1A shows that the expression of recombinant NOS enzymes allows a higher OD compared to bacteria expressing the empty vector (EV). The OD reached by bacteria expressing recombinant NOSs is 40 % higher than EV. The higher biomass produced by the expression of recombinant NOS in bacteria is shown in Supplemental Fig. S1 and reflected in the protein content and nitrate of cells (Supplemental Table S1), especially in the strain expressing SyNOS. The expression of OtNOS and SyNOS (119 kDa and 167 kDa respectively) was detected by immunoblot after 2 h of IPTG induction (Fig. 1B). Supplemental Fig. S2 shows that the effect of NOS expression on bacterial growth is completely abolished when bacteria are grown without IPTG addition. The increased growth rate of *E. coli* expressing SyNOS was reported previously (Correa-Aragunde et al. 2018), but was not tested for another NOS from photosynthetic microorganism like OtNOS.

**Figure 1.**
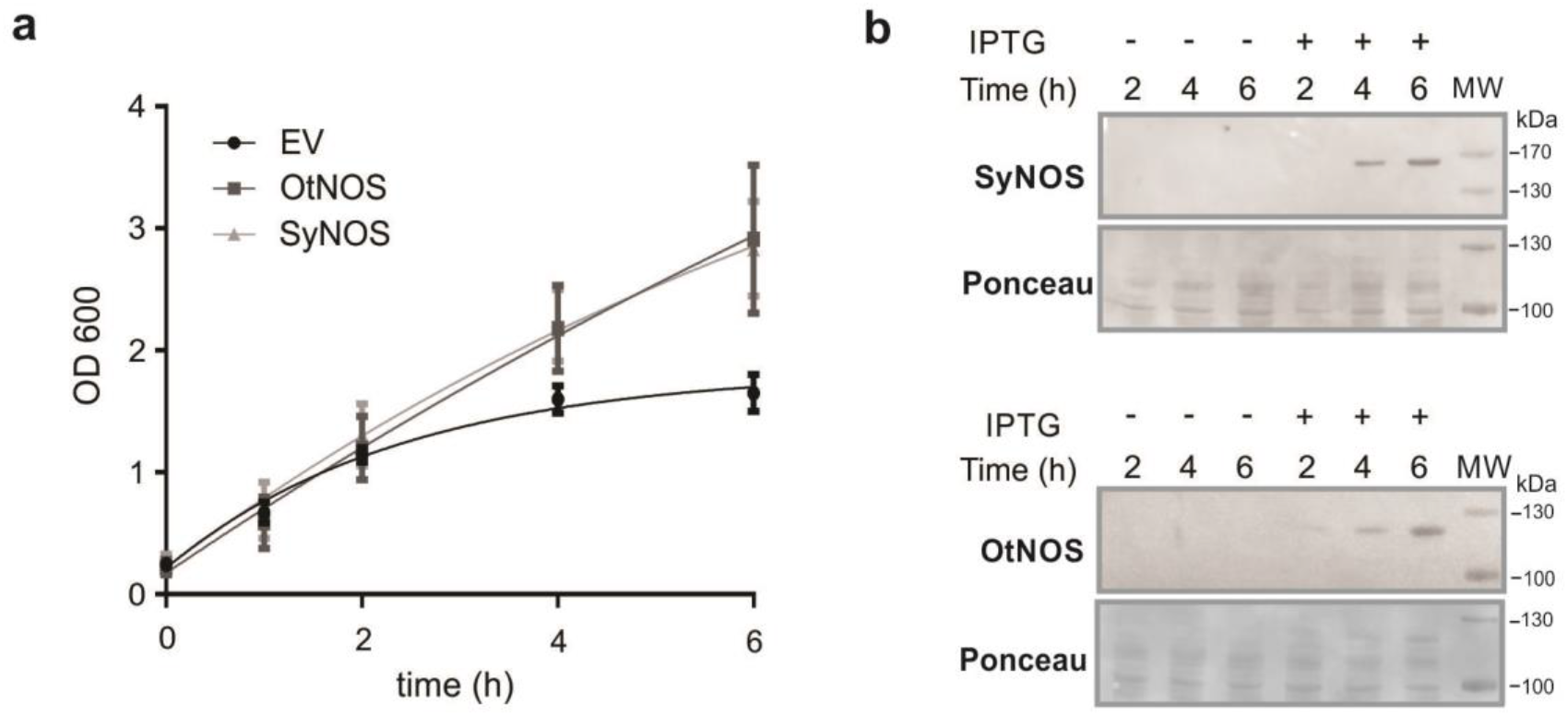
Growth curves of *E. coli* cultures that express OtNOS or SyNOS. **a** *E. coli* cultures transformed with the vector pET24a (EV), pET24a-OtNOS (OtNOS) or pET24a-SyNOS (SyNOS), growing in LB medium, were induced with 0.1 mM IPTG at OD=0.25. Growth was followed during different time points at OD 600. **b** Western blot showing the presence of OtNOS and SyNOS at the different time points with and without the addition of 0.1 mM IPTG. Full length blots are shown in Supplemental Fig. S3.

The bacterial NO dioxygenase gene (*hmp*) is considered a nitrosative stress sensor (Forrester and Foster 2012). HMP is capable of the conversion of NO to NO_3_^-^ in an O_2_-dependent manner. The expression of *hmp* was analyzed in *E. coli* after induction of OtNOS and SyNOS by quantitative PCR. The induction of *hmp* was detected after 4 h of OtNOS induction (Fig. 2). The OtNOS-dependent *hmp* induction reaches up to 6-fold respect to EV expressing bacteria. This result was not surprising since OtNOS appears to be an ultra-fast NO-releasing enzyme (Weisslocker-Schaetzel et al., 2017). SyNOS expression did not produce induction of the *hmp* gene at the same analyzed times (Fig. 2), suggesting that the NOD activity of the SyNOS-globin domain may be catalyzing the oxidation of NO to NO_3_^-^, thus these cells are not sensing nitrosative stress.

**Figure 2.**
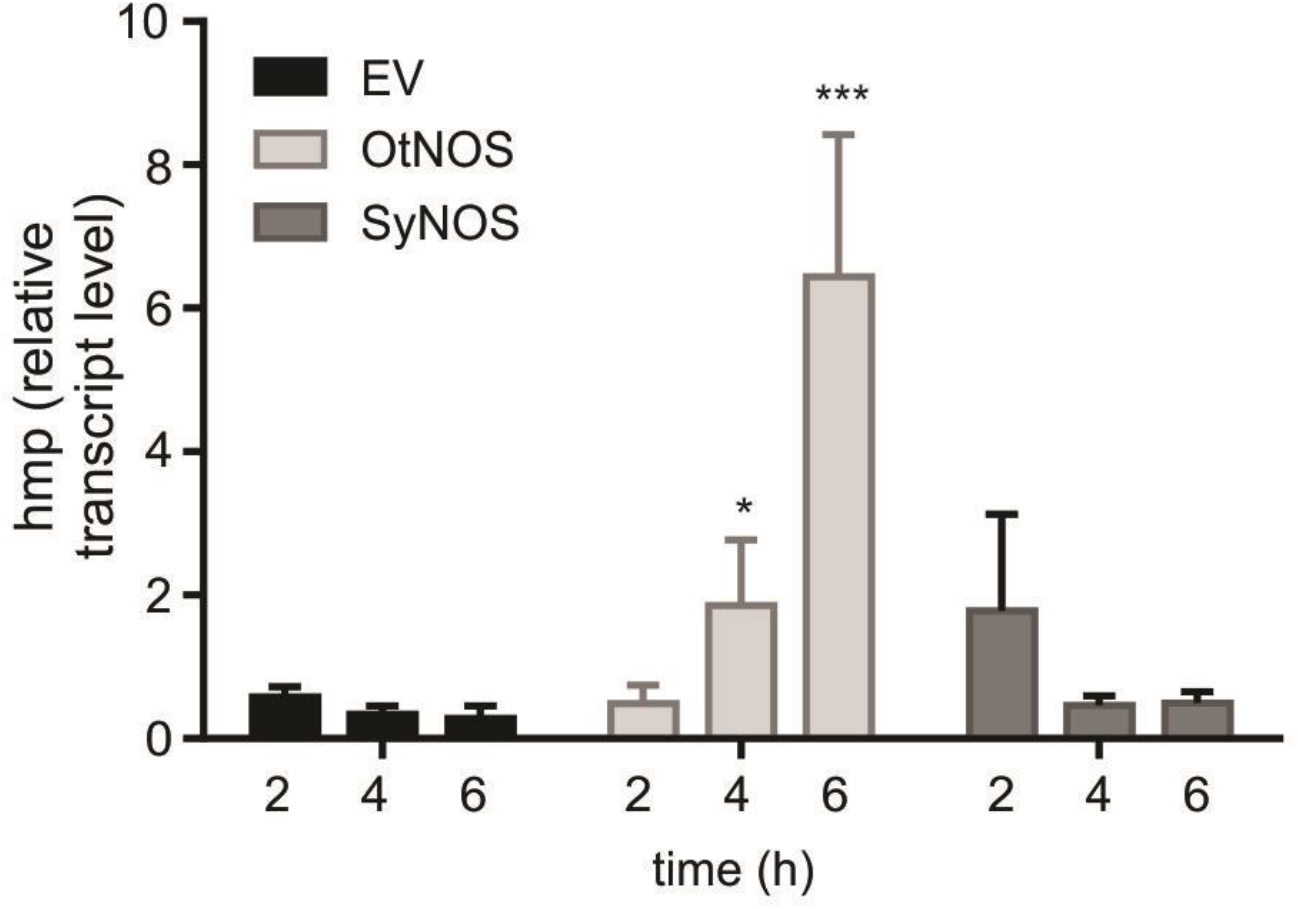
Expression of NO dioxygenase (*hmp*) in *E. coli* transformed with OtNOS or SyNOS. *E. coli* cultures (EV, OtNOS, SyNOS) were induced at OD = 0.25 with 0.1 mM IPTG. At the indicated time points, RNA was extracted, and cDNA synthesized. Expression of *hmp* was analyzed by qPCR. The expression of *hCat* transcript was used as housekeeping to relativize hmp levels. Values are means and SE of two independent experiments with one and two biological replicates (n=3). Asterisks means statistical difference respect to EV at the same time point (ANOVA, post hoc Dunnett method, *p < 0.1; ***p < 0.001).

### Effect of SyNOS and OtNOS expression on the *E. coli* growing in minimal medium with arginine as N source

Since NOS are arginine-degrading enzymes, we tested whether OtNOS expression in *E. coli* contributes to growth with arginine as a sole N source, as was previously evaluated with SyNOS (Correa-Aragunde et al., 2018). The hypothesis that we handle is that OtNOS may be able to produce NO and then *hmp* induction (Fig. 2) could catalyze the conversion of NO to NO_3_^-^. NOS expression was induced with IPTG addition for 1.5 h in LB medium and then an aliquot (100 μl) was transferred to 10 ml minimal medium containing 0.2 % (w/v) glucose and 0.2 % (w/v) arginine, maintaining the induction with 0.1 mM IPTG. Bacterial growth was followed by measuring OD 600. Figure A shows that OtNOS expressing bacteria, as well as SyNOS expressing bacteria, can grow with arginine as N source. At the times evaluated, it was negligible the growth of IPTG-induced EV bacteria, in contrast to another study (Picciano and Crane 2019). The maximum OD reached by OtNOS expressing bacteria is lower than SyNOS expressing ones, suggesting that OtNOS-HMP combined activity could be less effective than SyNOS containing the intrinsic NOD activity (Fig. 3). We also performed an experiment with a prior wash of cell pellets before the transfer to the minimal medium, as performed in Picciano and Crane (2019) (Supplemental Fig. S4). Some differences were observed such as delayed growth and less final OD reached by the strains, but again SyNOS-expressing strains generated the higher OD (Supplemental Fig. S4).

**Figure 3.**
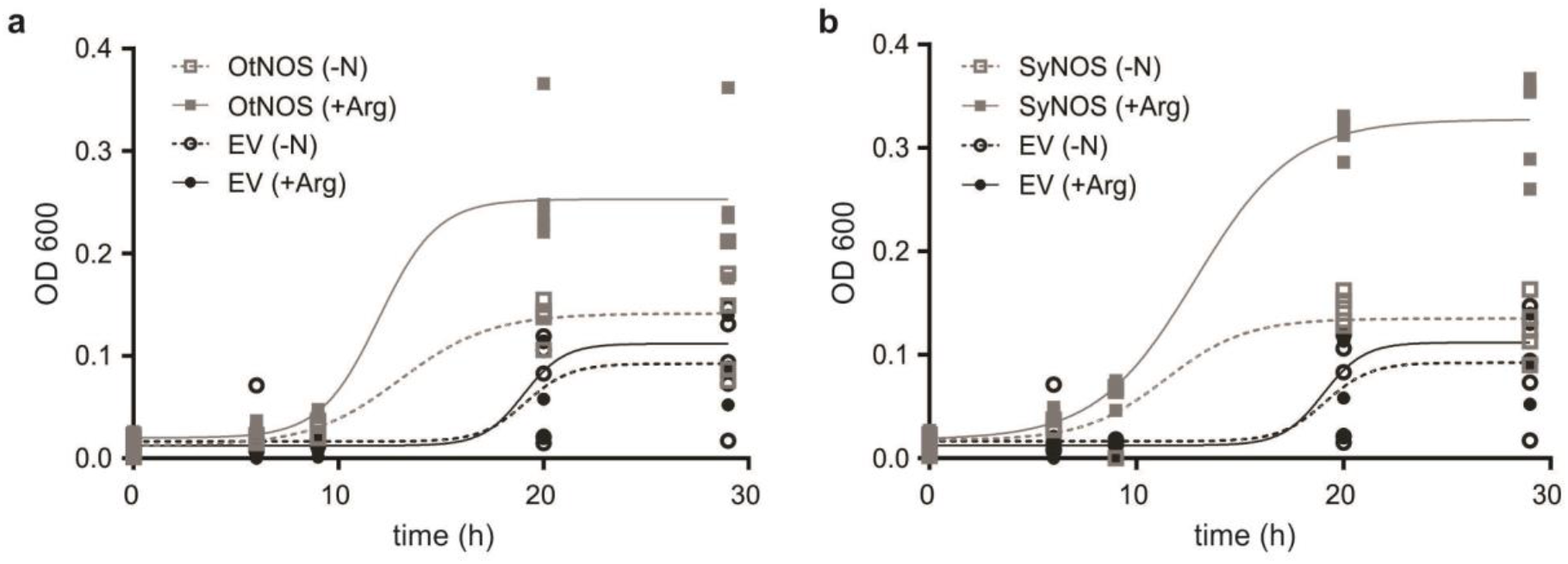
*E. coli* cultures that express OtNOS or SyNOS growing in minimal medium (MM) with or without arginine as a sole N source. *E. coli* EV, (**a**) OtNOS and (**b**) SyNOS cultures were induced with 0.1 mM IPTG for 1.5 h. Cultures were diluted 1/100 in MM containing 0.2 % (w/v) glucose without nitrogen source (-N) or with 0.2 % (w/v) arginine (+Arg), 0.1 mM IPTG and 50 μg/ml Kan. Growth was followed measuring OD 600 at different time points (n=5).

### The expression of recombinant OtNOS and SyNOS protects *E. coli* from nitrosative stress

Furthermore, we analyzed whether the expression of NOS in *E. coli* could protect bacteria from nitrosative stress. NOS expression was induced with IPTG addition for 1.5 h in LB medium and then was diluted to OD 0.2 maintaining IPTG induction and Kan selection. Nitrosative stress was induced by the addition of 1 mM sodium nitroprussiate (SNP). Growth was followed by OD 600 over 3 h with agitation to allow the adequate expression of the recombinant NOS proteins over the whole experiment. Figure 4 shows that NOS expressing culture achieved significantly higher μSNP /μno-stress ratios than the EV strain. The growth rates of *E. coli* cultures expressing either OtNOS or SyNOS were barely affected by nitrosative stress compared to EV cells, indicating that NOS expression aids cultures to counter SNP toxicity. OtNOS *E. coli* strain showed μSNP/μno-stress ratio of approximately 0.8, far more effective than SyNOS expression (μSNP/μno stress ratio 0.6). These results suggest that OtNOS expression may induce a priming effect since its expression induces the *hmp* gene (Fig. 2). On the other hand, partial SyNOS protection could come from SyNOS dioxygenase activity of the globin domain.

**Figure 4.**
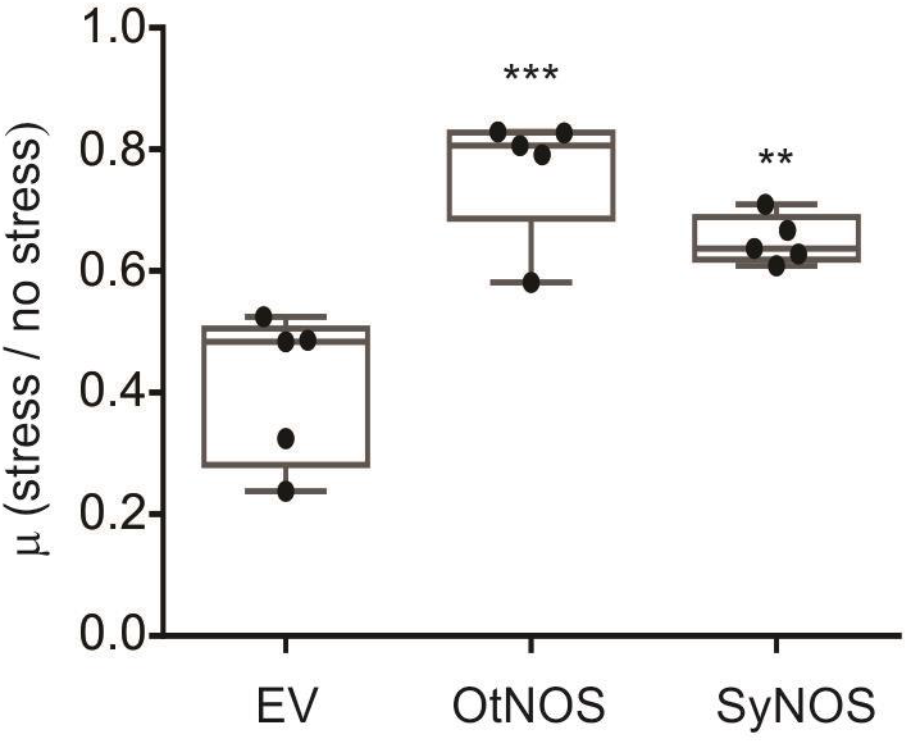
*E. coli* cultures expressing OtNOS and SyNOS are protected from nitrosative stress. *E. coli* cultures were induced for 1.5 h with IPTG 0.1 mM. Cultures were diluted in LB medium to OD 600 = 0.2. Nitrosative stress was triggered by addition of 1 mM sodium nitroprusside (SNP). Growth was measured by following OD 600 for 3 h. Data are expressed as the ratio of growth rate (μ) in the stress condition over non stress condition. Box plot from two independent experiments with two and three biological replicates (n=5). Asterisks indicate statistical differences compared to EV (ANOVA, post hoc Dunnett method, ** p < 0.01, *** p < 0.001). Representative growth curves of *E. coli* EV, OtNOS or SyNOS cultures subjected to nitrosative stress by SNP are shown in Supplemental Fig. S5.

### Analysis of putative pterins that could be operating as NOS cofactors in *E. coli*

Given the controversy about the type of pterin cofactor that NOS expressed in *E. coli* is using, and considering that: (i) BH4 is absent in this bacterium, which has not endogenous NOS and (ii) *in vitro* activity indicates that SyNOS does not utilize THF (Picciano and Crane 2019), we conducted an *in silico* study to shed some light on the matter. Both cofactors BH4 and THF share the pterin ring necessary for the electron donor process though its R side chain is different (Fig. 5A). However, some studies point out that L-tetrahydromonapterin (MH4), is one of the major pterins synthesized by *E. coli* (Ikemoto et al. 2002; Pribat et al. 2010). MH4 has a trihydroxypropyl as R side chain instead of the dihydroxypropyl chain present in BH4 (Fig. 5A). Taking into account that the R side chain of BH4 is not apparently involved in the electron donor process during NOS activity (Bird et al. 2002; Crane et al. 2010; Weisslocker-Schaetzel et al. 2017), this sole difference between both cofactors BH4 and MH4 allow us to postulate MH4 as the cofactor responsible for NOS activity in *E. coli*. This hypothesis is, in addition, supported by the fact that a structural isomer of MH4, the tetrahydroneopterin (NH4), is able to replace BH4 in mammal NOS catalysis under *in vitro* conditions (Presta et al. 1998; Gorren et al. 2001). Here, we generated tridimensional models for SyNOS and OtNOS using crystalized mammal NOS as template and utilized them in docking assay to assess BH4 and MH4 binding positions (Fig. 5B, C and D). Results indicate that almost all residues from the pterin pocket of mammal NOS are conserved in SyNOS and OtNOS with a remarkably similar structural alignment (RMSD 0.219Å and 0.481Å respectively). Only minor changes were detected in the position of key residues for the pterin binding between iNOS and SyNOS, that were also reported for OtNOS (Weisslocker-Schaetzel et al. 2017), suggesting that these differences may not affect NOS activity. I456 in iNOS is replaced by K391 in OtNOS (Fig. 5C) and G804 in SyNOS (Fig. 5D). Moreover, superimposed 3D structures show that the R723 guanidinium in SyNOS is displaced 1.8Å from R375 of iNOS (Supplemental Fig. S6).

**Figure 5.**
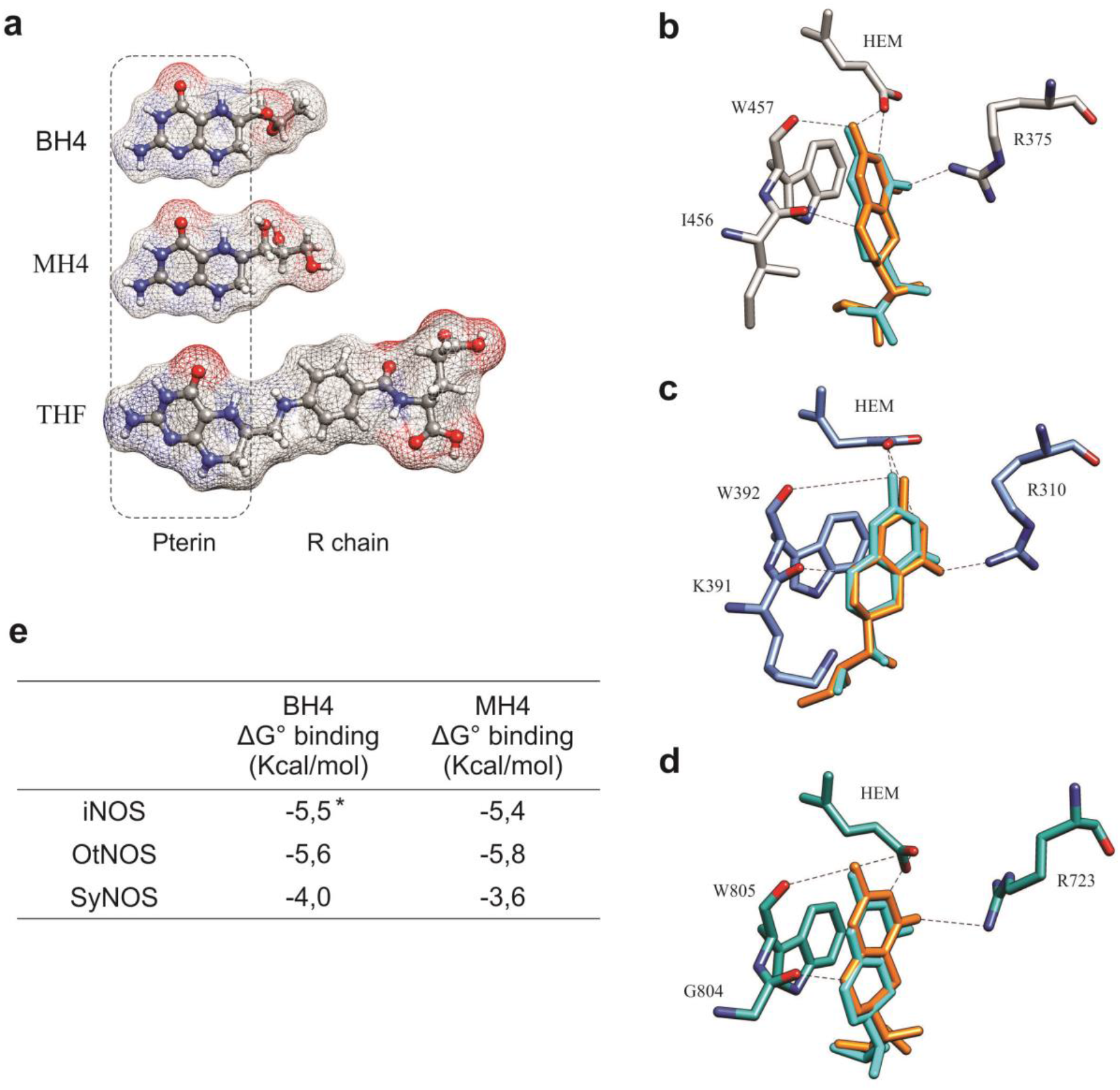
3D structures of the pterin binding residues showing H-bonds network in NOS enzymes. **a** Ligand structures in a ball and stick representation showing its electrostatic surface. Positive and negative charges are in blue and red, respectively. Positions of BH4 (cyan) and MH4 (orange) by molecular docking are shown in (**b**) iNOS 3D structure (PDB file 1M8D), (**c**) OtNOS and (**d**) SyNOS. **e** Ligand binding energy obtained after docking using AutoDock Vina software. Asterisk indicates energy obtained after self-docking.

In agreement with the structural changes in pterin binding residues described, molecular docking analysis revealed distinct positions for BH4 binding in OtNOS and SyNOS when it was compared with the crystalized position in iNOS (PDB 1M8D). Figures 5C and D show the position of BH4 in OtNOS and SyNOS that allow the binding (H-bonds and π-stacking) with the key residues necessary for cofactor stabilization and electron donation. Likewise, the possible binding position for MH4 cofactor performed by docking assay is like BH4 localization irrespective of the NOS analyzed (Fig. 5B, C and D). Delta free energy of BH4 and MH4 binding show similar negative values in all the NOS analyzed, though SyNOS showed a little less affinity due to the absence of residue interaction with pterin R chain (Fig. 5E). All at once, these evidences suggest that MH4 could bind in a similar position and affinity as BH4 and sustain NOS catalytic activity in *E. coli*.

## Discussion

In this work, we present evidence that heterologous expression of NOS proteins from photosynthetic microorganisms generates active enzymes in *E. coli*. The recombinant NOS expression promotes the growth in complete culture media and contributes to counteract nitrosative stress in *E. coli*. The involvement of bacterial NOS in cell proliferation and growth recovery after oxidative stress was previously reported (Gusarov and Nudler 2005; Shatalin et al. 2008; Gusarov et al. 2009). In *D. radiodurans*, UV irradiation induces upregulation of NOS gene, and NOS-derived NO upregulates obgE, a transcription factor involved in bacterial growth proliferation and stress response. Authors indicate that NO acts as a molecular signal for the regulation of growth in *D. radiodurans* (Patel et al. 2009). Likewise, ΔNOS strains of *B. subtilis* show a delayed growth response compared to the wt. It seems that NO is required for maintaining normal cell growth rate in *B. subtilis*, mainly at the entering of the stationary phase (Gusarov and Nudler 2005).

It is well known that IPTG addition has a negative impact on bacterial cell growth and compromises the biochemistry and physiology of the strain used for recombinant expression (Kosinski et al. 1992; Malakar and Venkatesh 2012; Lecina et al. 2013; Dvorak et al. 2015). IPTG achieves both, high levels of foreign gene expression and a high cell density and, at the same time, it is difficult for the host bacteria because of: (i) the accumulation of toxic waste products (i.e. acetic acid), (ii) the consumption of precursors to allow the synthesis of foreign proteins and selection markers and (iii) the replication of plasmid vectors. In this context, IPTG addition affects the final OD reached by the EV strain while OtNOS and SyNOS expression can partially overcome this restriction when grown in complete LB medium (Fig. 1 and Supplemental Fig. S1). This result suggests that NOS expression could be applied as a tool to keep the potential of bacterial growth during the production of recombinant proteins of commercial interest.

Picciano and Crane (2019) inquired about the nature of the pterin cofactor needed for SyNOS activity in *E. coli* since this bacterium does not have sepiapterin reductase involved in the synthesis of BH4 and, additionally, SyNOS is not active with THF at least in *in vitro* assays. THF is a molecule that contains a bulkier R side chain (glutamyl p-amino benzoic acid, pABA) than BH4 and MH4 (Fig. 5A), needing more physical space in the NOS pterin pocket. Also, this pABA side chain has to be surrounded by residues that provide electrostatic stability to preserve its binding for electron donation. Crane et al. (2010) established that mammals NOS are not structurally able to bind THF. Although this cofactor seems to be functional for OtNOS (Foresi et al. 2015), Weisslocker-Schaetzel et al. (2017) have described the structural disadvantages that THF could have for its binding. In contrast, bacterial NOSs (i.e. *B. subtilis, D. radiodurans*) have a more open pterin pocket, allowing THF to bind and act as a redox cofactor (Adak et al. 2002b; Adak et al. 2002a). Nevertheless, there still remains some doubts about the native NOS cofactor in bacteria, since THF fails to activate NOS from *Staphylococcus aureus* (Bird et al. 2002). In the case of SyNOS, where its dimerization structure is unknown, and the whole enzyme folding with its globin domain has not been established, further analysis is needed to determine the specific disability for THF binding.

Curiously, it was demonstrated that the oxygenase domain of rat nNOS is functionally active in *E. coli*, although this NOS also exclusively uses BH4 instead of THF (Gusarov et al. 2008). Authors observed a higher NO_2_^-^ and NO_3_^-^ production in *E. coli* cells expressing a rat nNOS or *B. subtilis* NOS when compared to BL21 control bacteria. However, the authors did not refer to which pterin cofactor nNOS may be using (Gusarov et al. 2008). Gorren et al. (2001) analyzed which of many pterins are able to function as human NOS cofactors due to their redox properties. Interestingly, they have shown that NH_4_, a structural isomer of MH4, could support NOS activity. *E. coli* synthesizes considerable amounts of MH4 by the action of two characterized enzymes called FolX (neopterin epimerase) and FolM (monapterin reductase) (Pribat et al. 2010). Also oxidized monapterin was found to be excreted in the bacterial culture medium, however the role of the excreted pterin by bacteria is still unknown (Wachter et al. 1980). In *Pseudomonas aeruginosa*, MH4 seems to act as the cofactor of the enzyme phenylalanine hydroxylase (Pribat et al. 2010), a BH4-dependent enzyme in animals. In agreement with this, the analysis of the recombinant expression of a mouse BH4-dependent tyrosine hydroxylase in *E. coli*, only showed activity in wt strains, whereas in mutant strains (*folX* and *folM*) lacking MH4 synthesis route, no activity was detected (Satoh et al. 2012). These results strongly indicate that MH4 can replace BH4 in a human BH4-dependent enzyme.

When the genome of *Synechococcus* PCC was analyzed to find a gene coding for a sepiapterin reductase (SPR), one sequence revealed a 30% similarity, next to the gene encoding GTP cyclohydrolase I (GTPCH I). This sequence encodes for a short-chain dehydrogenase (SDR) domain and authors speculated that this strain may synthesize BH4 (Picciano and Crane 2019). However, the SDR family is a large family of oxidoreductases (Kavanagh et al. 2008) and this sequence lacks the specific motif of the pteridine reductases (an Arg in place of the Gly at position 6 in the TGX_3_RXG motif) (Labine et al. 2020). Thereby, it remains to be elucidated whether *Synechococcus* PCC 7335 can synthesize BH4. Here we show that the expression of both SyNOS and OtNOS slightly increases the *E. coli* growth in minimal medium, and this contrast to the result showed by Picciano and Crane (2019). We postulate that the addition of IPTG to maintain NOS expression in our experiments described in this article and those previously reported (Correa-Aragunde et al, 2018), is the main difference compared to experiments performed in Crane’s lab., and that it could be responsible for the differing results.

In summary, our results show that recently described NOSs from photosynthetic organisms increase the growth and confers oxidative stress tolerance in *E. coli*. These properties might be interesting to apply in biotechnological uses, such as production of polymers, secondary metabolites and antibiotics in *E. coli*. Besides, the analysis allows us to firmly postulate MH4 as a pterin cofactor in *E. coli* to sustain NOS activity. These results open new windows to discuss about the versatility of pterin cofactors working in NOSs.

## AUTHOR CONTRIBUTIONS

NC-A, FDC and N.F designed and performed all experiments. A.N conducted statistical and ligand docking analysis. NC-A and AN wrote the manuscript. NC-A and LL analyzed the data and initiated and supervised the project.

## ACKNOWLEDGMENTS

We acknowledge Mar del Plata University and the Consejo Nacional de Investigaciones Científicas y Técnicas (CONICET). FC and AN are PhD fellows and NF, LL, and NC-A are members of the research staff from CONICET, Argentina.

## ETHICAL STATEMENT

This article does not contain any studies with human participants or animals performed by any of the authors.

## Data availability statement

The data that support the findings of this study are available from the corresponding author upon reasonable request.

