## Supplemental File for "Nitric oxide synthases from photosynthetic organisms improve growth and confer nitrosative stress tolerance in *E. coli*. Insights on the pterin cofactor"

**Journal name: Applied Microbiology and Biotechnology**

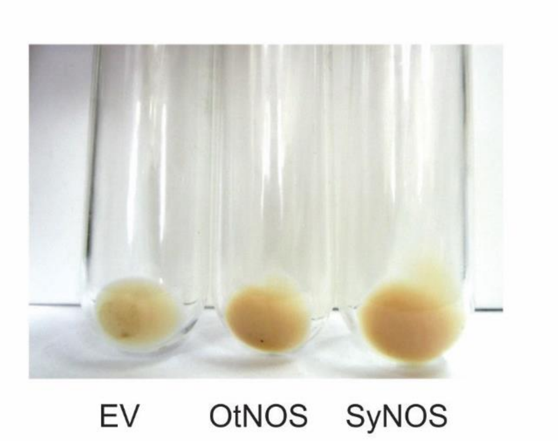


Supplemental Fig. S1. Cells pelleted from liquid cultures of *Escherichia coli* that express the empty vector (EV), or OtNOS or SyNOS recombinant proteins. *E. coli* cultures transformed with the vector pET24a (EV), pET24a-OtNOS (OtNOS) or pET24a-SyNOS (SyNOS) growing in LB medium were induced with IPTG 0.1 mM at OD=0.25 and growth for 6 h at 30 ºC. Twenty ml of cell cultures were pelleted by centrifugation at 5,000 rpm during 10 min.

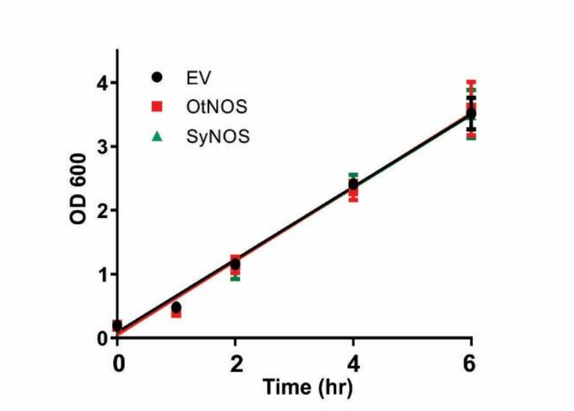


Supplemental Fig. S2. Growth curve of *E. coli* cultures that express the empty vector (EV), OtNOS or SyNOS proteins without the addition of IPTG. Growth was followed by measuring OD 600 at different time points. Data are means and SE from at least 3 independent experiments.

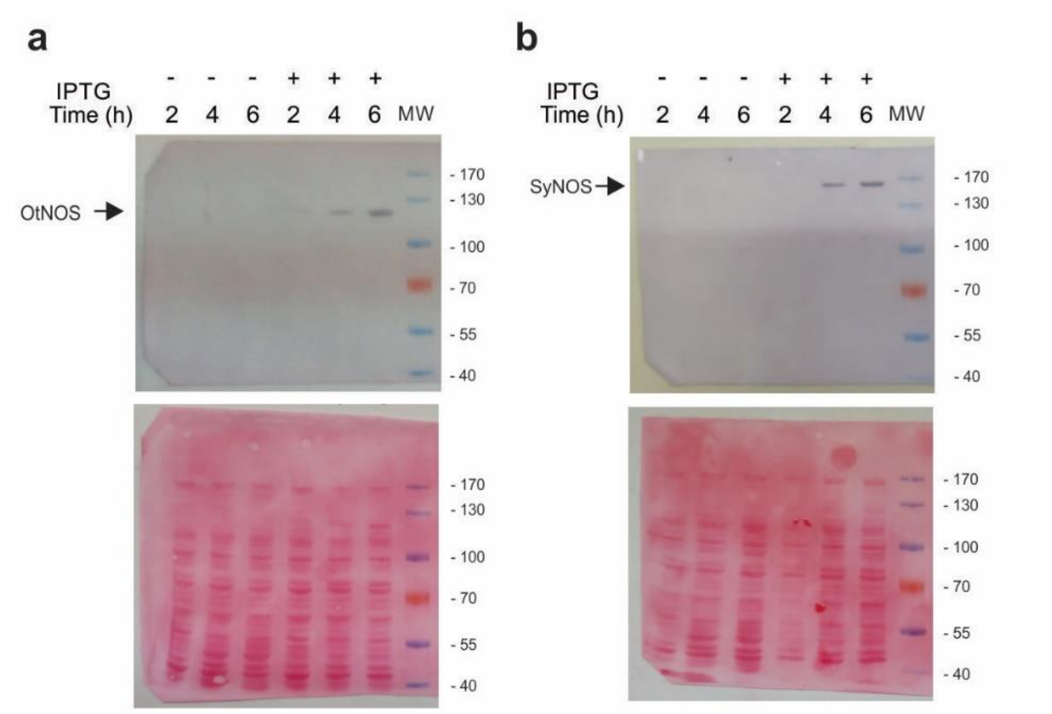


Supplemental Fig. S3. Full length western blot (upper panel) and Ponceau staining (bottom panel) showing the presence of the recombinant proteins OtNOS (**a**) and SyNOS (**b**) at the different time points with and without the addition of IPTG 0.1 mM. MW: molecular weight markers.

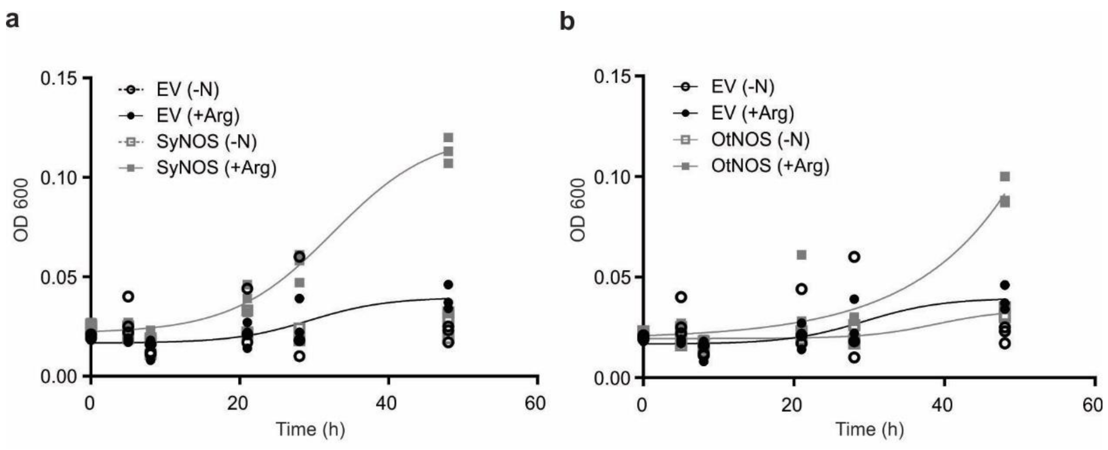


Supplemental Fig. S4. Growth of *E. coli* cultures that express OtNOS or SyNOS in minimal medium (MM) with arginine as a sole N source. E. coli EV, (**a**) SyNOS and (**b**) OtNOS cultures were induced with 0.1 mM IPTG for 1.5 h. Aliquots were centrifuged and washed three times with MM without N and then resuspended in minimal media containing 0.2 % (w/v) glucose without nitrogen source (-N) or with 0.2 % (w/v) glucose plus 0.2 % (w/v) arginine (+Arg). Growth was followed by measuring OD 600 at different time points.

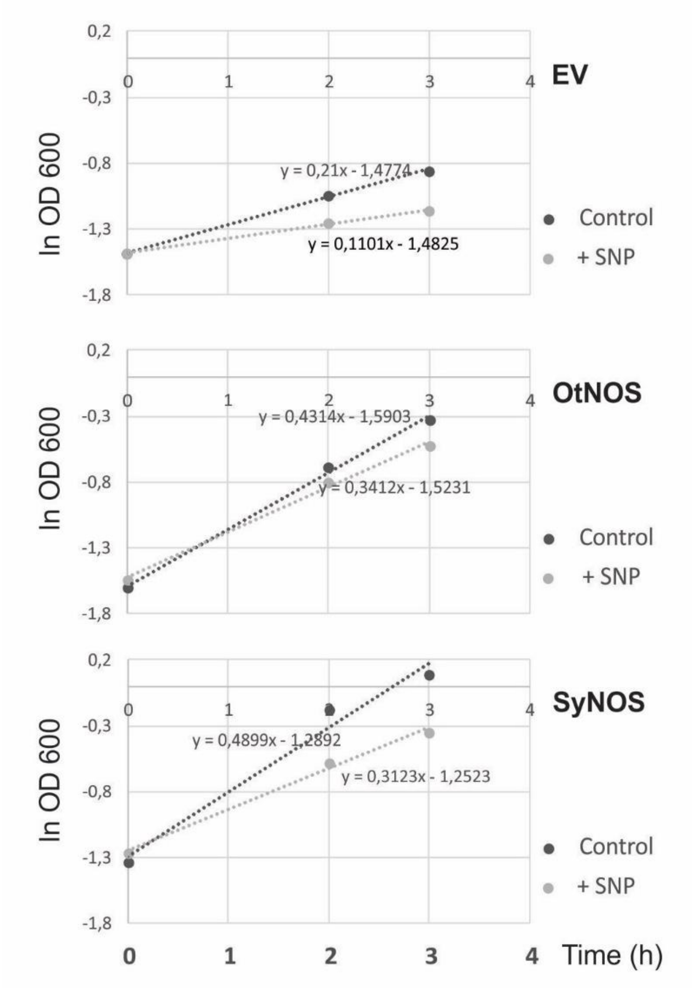


Supplemental Fig. S5. Growth curves of *E. coli* cultures that express the empty vector (EV), OtNOS or SyNOS under nitrosative stress induced by SNP. *E. coli* cultures were induced for 1.5 h with 0.1 mM IPTG. Cultures were diluted in LB medium to OD 600 ~ 0.2. Nitrosative stress was triggered by addition of 1 mM SNP. Growth rates (µ) were calculated by applying linear regression analysis on the natural logarithm (ln) of Abs 600 values versus time of culture as described by Frey et al. (2002). Representative growth response curve measured by following OD 600 for 3 h.

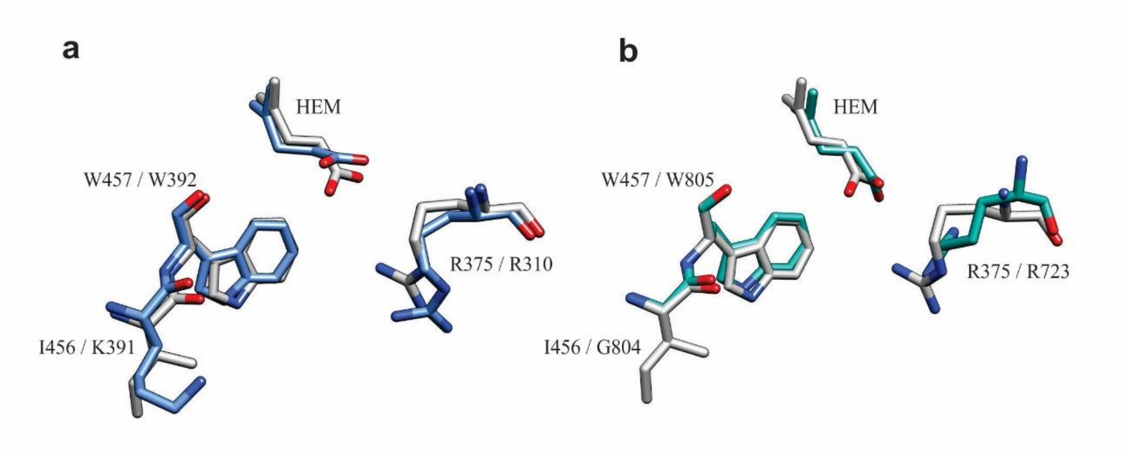


Supplemental Fig. S6. Spatial positions of pterin binding residues in OtNOS and SyNOS. Residues in licorice representation from OtNOS (blue) (**a**) or SyNOS (green) (**b**) were superimposed with iNOS residues (grey) in a 3D scenario. Residues are indicated as iNOS / OtNOS or SyNOS respectively.

Supplemental Table S1. Protein, nitrate and nitrite content in *E. coli* cultures that express OtNOS or SyNOS

| | Protein (µg/µl) | Nitrite (µM) | Nitrate (mM) |
|---|---|---|---|
| EV | 0.310 ± 0.07 | 33.61 ± 4.98 | 3.53 ± 0.40 |
| OtNOS | 0.616 ± 0.09 | 20.02 ± 2.25 | 3.45 ± 0.20 |
| SyNOS | 1.170 ± 0.17*** | 30.29 ± 4.12 | 6.02 ± 0.11*** |

*E. coli* EV, SyNOS and OtNOS were induced in 3 ml culture by the addition of 1 mM IPTG for 6 hs. Samples were collected by centrifugation at 5000 rpm for 5 min and resuspended in 200 µl PBS 1X. Protein concentration was analyzed with Bradford reagent; nitrite content was evaluated with the Griess method and nitrate by the technique reported by Cataldo et al. (1975). Data are means ± SE of 5 biological replicates. Asterisks indicate statistical differences compared to EV (ANOVA, post hoc Dunnett method, *** $p < 0.001$)
